# Whole genome sequencing reveals a novel *Renibacterium salmoninarum* lineage and suggests geographic endemism combined with anthropogenic spread in the North-East Atlantic area

**DOI:** 10.1101/2025.08.18.670871

**Authors:** Alejandra Villamil-Alonso, Torfinn Moldal, Cathrine Arnason Bøe, Birkir Thor Bragason, Hanne Nilsen, Anne Berit Olsen, Åse Helen Garseth, Snorre Gulla, Niccolò Vendramin, Lone Madsen, Argelia Cuenca, Karin Lagesen

## Abstract

*Renibacterium salmoninarum* is the causative agent of bacterial kidney disease (BKD) and has been isolated from northern European salmonid farms since the 1960s. The bacterium has been detected only sporadically in Norway during the last decades, but the country experienced several geographically spread outbreaks since December 2022. The phylogenomic relationships of *R. salmoninarum* isolates associated with the epizootics ongoing on the West Coast and in Mid-Norway were explored using a whole genome sequencing approach. A broad overview of the phylogeography of this pathogen was gained through sequencing and analysis of a collection of isolates from Norway (n=67), Iceland (n=12), Denmark (n=12), and the Faroe Islands (n=1), along with a collection of 109 publicly available sequences. We identified two distinct contemporary clades of *R. salmoninarum* causing BKD in Norway in the period 2022-2024. Both clades belong to the expanding, aquaculture-associated Lineage 1. The epidemiological picture appears consistent with contemporary aquaculture operations, raising questions on the effectiveness or practice of current biosecurity practices towards *R. salmoninarum*. Our work also describes a hitherto undescribed lineage (Lineage 3), predominantly from Iceland, where BKD is considered endemic. The detection of endemic reservoirs of *R. salmoninarum* in European water systems underscores the potential for ongoing pathogen circulation independent of acute outbreaks. This finding emphasizes the importance of further investigation of the mechanisms of pathogen persistence, particularly within environments related to aquaculture, where chronic infection reservoirs could compromise disease management and biosecurity.

**IMPORTANCE:** *Renibacterium salmoninarum*, the causative agent of bacterial kidney disease (BKD), is a chronic salmonid pathogen that often presents subclinically, complicating detection and control. Effective management of BKD is therefore crucial for sustainable aquaculture and wild fish conservation. Recognizing the need for comprehensive sampling, previous studies have underscored the importance of a global isolate collection to better understand the epidemiology of the pathogen. While the slow genetic evolution and highly conserved genome of the bacterium have reduced the resolution of conventional typing methods in the past, applying whole genome sequencing has enabled its molecular tracing at the outbreak level. Here, we present the largest phylogenomic analysis of *R. salmoninarum* to date, including sequences from Denmark, Iceland, and the Faroe Islands for the first time. This expanded dataset elucidates past and present movements of the pathogen in European waters, providing actionable insights to support development of targeted BKD management and biosecurity strategies.

## INTRODUCTION

Global aquaculture has undergone substantial growth, with aquaculture production of animal species surpassing capture fisheries for the first time in 2022 (1). This rapid expansion, coupled with the reliance of the industry on surrounding ecosystems such as the quality of natural water bodies for water exchange, waste management, and the maintenance of optimal rearing environments, has heightened its vulnerability to a range of environmental stressors, including the introduction and persistence of pathogens (2). Among these, *Renibacterium salmoninarum* represents a critical threat. This Gram-positive, slow growing, non-spore forming, facultative intracellular coccobacillus (3) belongs to the high GC content group (4), and is the etiological agent of bacterial kidney disease (BKD), which affects wild and farmed salmonid fish in fresh and seawater worldwide (5). The bacterium is capable of both horizontal (6, 7) and vertical transmission (8, 9), facilitating its persistence in affected fish populations (10). BKD typically manifests as a chronic, granulomatous infection with slow progression and frequent subclinical presentation (3, 7). Wild salmonids as well as other marine and freshwater fish species have been identified as reservoirs for the bacterium (11), and a lifelong carrier state has been proposed (12). The survival of *R. salmoninarum* within chronically infected hosts, coupled with the ability to transmit vertically, enhances its persistence and complicates disease control (10, 12). Moreover, current diagnostic methods may fail to detect subclinical BKD infections, further hindering effective disease management (12–14). The presence of BKD is not a recent phenomenon in Europe (15–17) and the bacterium has been isolated from European salmonids since the 1960s, including regions like Denmark, Iceland, and Norway (17–20) after the first documented isolation of the pathogen form wild Atlantic salmon (*Salmo salar*) in the River Dee during the early 1960s. Prevalence, morbidity, and mortality rates vary widely among fish species and with individual outbreaks, often with minor or no overt clinical signs (16, 21, 22) even when infection prevalence is high (17). However, severe cases of BKD have been documented where losses in cultured salmonids can range from 5-40% (23) and with some outbreaks leading to losses of up 133 tones per farm (11).

Norway is the largest salmonid aquaculture producer in the world, contributing 1-2% of the global aquaculture production, primarily Atlantic salmon (1, 2). It also sustains approximately 450 of the world’s 2,000 wild Atlantic salmon populations, distributed across rivers from Skagerrak in the south to the Barents Sea in the north (24). The dramatic decline in wild Atlantic salmon populations has prompted several conservation and protection measures, including expansion of the national gene bank program (25). *R. salmoninarum* is present in Norwegian waters (26) and was first detected in Norway in 1980 (19), and BKD has been notifiable since 1969. The disease is currently classified in national category F, which dictates immediate notification to the Norwegian Food Safety Authority upon suspicion based on clinical or laboratory findings (27). The incidence of the disease peaked in 1990 with 62 affected sites and salmon from the rivers Eidselva and Valldalselva diagnosed with BKD (Håstein, NVI, pers. com.). Following the implementation of enhanced biosecurity measures in 1991, including the legal requirement for mandatory health checks of broodfish, BKD detections declined markedly (18). During the 1990s, these measures were implemented in stock enhancement hatcheries and gene banks. Since 2017, the Norwegian coastline has for regulatory purposes been divided into 13 production areas (PA), ranging from PA1 in the south to PA13 in the northeast, bordering Russia (28); see Fig. S1. Cases of BKD (i.e. confirmed presence of *R. salmoninarum* at a specific farming site during the same production cycle) have been detected along the Norwegian coastline since 1984. For this study, isolates from seven of these production areas were analyzed (Fig. 1). The mandatory annual screening of more than 600 wild broodfish has provided valuable insight into the presence of *R. salmoninarum* in wild salmonids in Norway. While surveillance from 1992-1996 detected only six cases among 4,048 examined salmonids (0.15%) (29), recent decades have seen infection with *R. salmoninarum* in four rivers (Ekso, Lærdal, Daleelva, Vosso) (30), three of which are found in the same fjord system in the western part of Norway (PA4). A recent increase in BKD cases was registered in 20 aquaculture farming sites in PA4, 5, and 6 between December 2022 and the end of 2024, with one site affected twice (two cases) (31). As a result, an official surveillance program in wild salmonids has been put in place in Norway (30) and isolates have been collected from the cases ongoing since the start of the epizootics.

**Fig 1.**
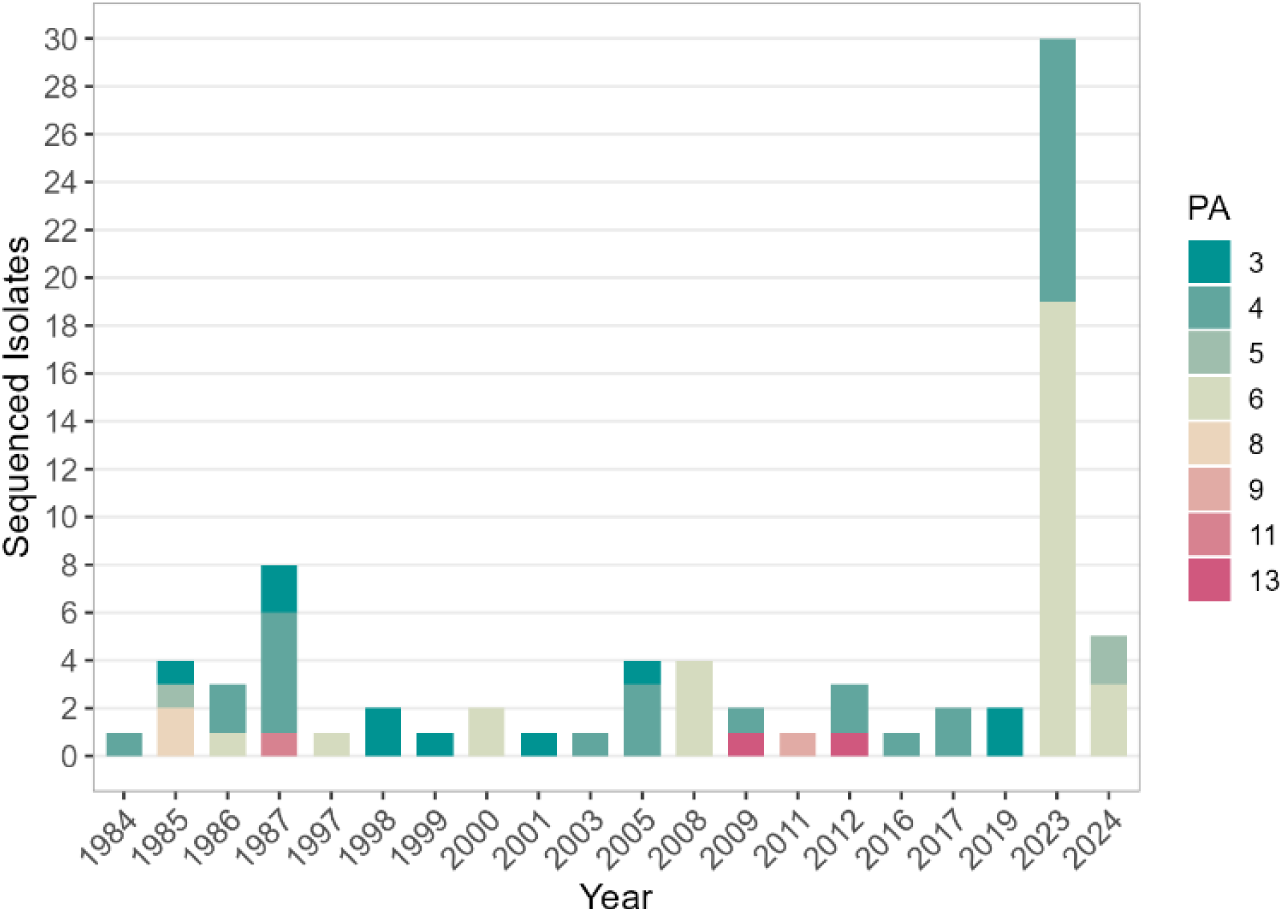
Temporal and spatial distributions of sequenced *Renibacterium salmoninarum* isolates in Norway, 1984-2024. The histogram displays the annual number of *R. salmoninarum* collected and sequenced from Norwegian cases of BKD, grouped by production area (PA). Each color represents a different PA, as indicated in the legend. Data shows all sequenced isolates from affected sites since 1984.

The first report of BKD in Iceland dates from 1968 and was associated with fish farming (21). The disease has been detected on a regular basis in connection with fish farming activities in the country since the 1980s (21, 32, 33). Routine monitoring of *R. salmoninarum* in wild salmon broodfish from Icelandic rivers has been performed annually since 1986 (21), with *R. salmoninarum* consistently detected, typically in less than 5% of salmonids, although a prevalence above 20% was observed in 2008-2009 (21). This, along with results from a survey performed by Jónsdóttir et al. that showed a widespread presence of *R. salmoninarum* infection in wild Arctic char (*Salvelinus alpinus*) and brown trout (*Salmo trutta*) populations in Iceland (17), indicates that *R. salmoninarum* is endemic and common in the three salmonid species that inhabit Icelandic water systems.

In Denmark, BKD was found for the first time in 1997, when *R. salmoninarum* was isolated during disease investigation of clinically affected rainbow trout (*Oncorhynchus mykiss*) in a farm sourcing water from the river system of Skjern Å, southwest Jutland (20). As the pathogen soon thereafter was detected in five other farms (among them a hatchery) in connection with the same river system, an eradication program for the disease was cancelled (20) and, instead, *R. salmoninarum* became and is still part of a national control program in Denmark. Thanks to the implementation of strict biosecurity measures and regular targeted surveillance, 14 compartments are currently declared officially free from the disease (34).

Accurate strain identification helps to trace the present and past movements of bacteria and greatly benefits from the use of highly discriminatory approaches, such as whole genome sequencing (WGS) (35). This is particularly valuable to study *R. salmoninarum*, whose genome (∼3.15 Mbp), like many intracellular bacteria, shows signs of reductive evolution (36) and constrained genetic intra-species variation. *R. salmoninarum* was the first aquaculture pathogen for which a global collection of isolates was investigated using next generation sequencing platforms, enabling high-resolution discrimination based on single nucleotide polymorphism (SNP) data across broad geographical, temporal, and ecological ranges. Brynildsrud et al. described two main clades with contrasting origins, designated Lineage 1 and Lineage 2 (26). While Lineage 2 was restricted to Scotland and Norway and associated with the historical endemic presence of the bacterium in European waters, Lineage 1 was represented in all of the countries studied, and its emergence and expansion was linked with increased anthropological activities in relation to aquaculture (26). Aquaculture operations as a driver for the spread of the *R. salmoninarum* from Lineage 1 was further supported by Bayliss et al., who studied a collection of Chilean isolates in the context of the global phylogeny of the bacterium (37). They found all Chilean isolates to belong to Lineage 1 and suggested that the introductions and local spread of the bacterium within the country were linked to changes in the aquaculture industry and subsequent aquaculture operations (37).

In this study, we investigated the phylogenomic relationships among *R. salmoninarum* isolates cultivated from BKD cases detected since December 2022 in PA4, PA5, and PA6 on the West Coast and in Mid-Norway. We aimed to clarify the epidemiological links behind these epizootics and to explore potential patterns of transmission of *R. salmoninarum*. To do so, we have performed the largest WGS phylogenomic analysis of *R. salmoninarum* to date. It includes 109 publicly available isolates (26, 37) for which raw sequencing reads were retrieved, and 92 newly sequenced isolates from Norway (n=67), Iceland (n=12), Denmark (n=12), and the Faroe Islands (n=1). This has allowed us to get a broader overview of the evolutionary relationships of the bacterium within the North-East Atlantic region, and to shed light on the phylogenomic links between recent cases.

## MATERIALS AND METHODS

### Collection of isolates

A total of 92 bacterial isolates were sequenced for the present study (Table 1, Table S1). All had been previously identified as *R. salmoninarum* by routine diagnostic methods from farmed salmonid fish within Norway, Denmark, Iceland, and the Faroe Islands. Most of the Norwegian isolates were obtained through laboratory diagnostics at the Norwegian Veterinary Institute. Samples from suspected cases were collected either by fish health services or Food Safety Authority personnel and submitted to the laboratory for disease investigation. Some of the samples from wild fish were obtained during surveillance programs and mandatory testing of broodfish, i.e. BKD was not suspected. The Icelandic samples were obtained through diagnostic activities at The Institute for Experimental Pathology at Keldur, University of Iceland, and the Danish samples were collected during voluntary surveillance and screened through diagnostic activities at the National Institute of Aquatic Resources (Kongens Lyngby, Denmark).

**Table 1.** Summary metadata of the 92 *Renibacterium salmoninarum* isolates sequenced for this study. The table presents the number of isolates sequenced from each country, categorized by host species and indicating the year range of isolation.

| Country | Number of isolates | Host species |  |  |  | Year range |
| --- | --- | --- | --- | --- | --- | --- |
|  |  | *S. salar* | *O. mykiss* | *S. trutta* | *S. alpinus* |  |
| Norway | 67 | 59 | 7 | 1 | - | 1985–2024 |
| Denmark | 12 | - | 12 | - | - | 2017–2023 |
| Faroe Islands | 1 | 1 | - | - | - | 1992 |
| Iceland* | 12 | 8 | 1 | - | 1 | 1987–2012 |

\*For Iceland, host species information is missing for two of the isolates. See Table S1 for complete metadata, including accession numbers.

### Bacterial culture and DNA extraction

The isolates were grown on SKDM and/or KDM agar media (38) and incubated at 15°C for single colony isolation at either the Norwegian Veterinary Institute (Bergen, Norway), The National Institute for Aquatic Resources (Kongens Lyngby, Denmark), or The Institute for Experimental Pathology at Keldur, University of Iceland (Keldur, Iceland). In Norway, suspected colonies were identified with MALDI-TOF MS (matrix-assisted laser desorption/ionization time of flight mass spectrometry) (Bruker®) before submitted to whole genome sequencing. Duplicate strains were stored at -80°C. All the Norwegian, Danish, and Faroese isolates were sequenced at the Norwegian Veterinary Institute (Ås, Norway).

Bacterial cell lysis was performed by overnight incubation with lysozyme (Merck, 62970) (20 mg/ml in 2mM EDTA, 1.2% Triton, 20mM Tris-HCl pH 8.0) at 37°C and 300 rpm. After pretreatment DNA extraction was performed using the QIAamp DNA Mini kit (QIAGEN, Q51306) according to the manufactureŕs instructions. For the Icelandic isolates genomic DNA isolation was performed using the GeneJet Genomic DNA Purification Kit (Thermo Scientific, #K0721) at The Institute for experimental pathology at Keldur, University of Iceland.

### Whole genome sequencing, assembly, and annotation

Genomic DNA quantification was performed using a Qubit 2.0 Fluorometer (Life Technologies, Thermo Scientific) and libraries were constructed from an input of ∼200 ng gDNA using Illumina DNA Prep (Illumina, 20060059). The libraries were sequenced using the MiSeq Reagent Kit with v3 chemistry and 600 cycles (Illumina, MS-102-3003) to obtain 2 x 300 bp paired end reads on an Illumina MiSeq. All steps were conducted following the manufacturers’ recommendations. For the Icelandic isolates, genomic DNA was sent to BGI Genomics which performed library construction and whole genome sequencing on the DNBSEQ platform (150 bp paired end reads).

Following sequencing, subsequent quality assessment, trimming, and assembly were conducted using the draft genome assembly track from the Assemblage pipeline for prokaryotic organisms (39). Briefly, paired-end Ilumina short reads were quality checked with FastQC v0.12.1 and MultiQC v1.14 (40, 41), trimmed with Trim-Galore v0.6.10 (Babraham Bioinformatics - Trim Galore!), and screened for contamination with Kraken v2.1.3 (42). To control sequencing depth, random subsampling of the read files was performed using rasusa v0.8.0 (43), followed by a second quality check on the output. Assembly was conducted using Unicycler v0.5.0 with default parameters and --genome_size set to 3,155,250 based on the *R. salmoninarum* ATCC 33209 reference strain (36, 44). Coverage was calculated by mapping reads to their respective assemblies using BWA v0.7.8, SamTools v1.3.1, and BedTools v2.31.0. Quality metrics for each individual assembly were obtained using Quast v5.2.0 (45). Additionally, short reads from a previously published set of 109 *R. salmoninarum* isolates were retrieved (26, 37). Fastq files were downloaded using SRA-Toolkit v3.0.3 on June 26^th^, 2024, and assembled as described above.

### Phylogenetic analysis

Whole genome similarity for the 201 assembled genomes was determined using the FastANI track from the pipeline ALPPACA v2.2.1 (46) which calculates average nucleotide identity (ANI) (<u>ParBLiSS</u>/<u>FastANI,</u> v1.33). Based on the obtained ANI values (pairwise ANI ≤99.8%), the “core genome” track was chosen for subsequent genome analyses. In short, ParSNP v1.6.1 was used to generate a colinear whole genome multiple alignment. Gubbins v3.1.6 was used to detect eventual recombination (47) and suspected recombinant areas were masked with maskrc-svg v0.5 to minimize the impact of recombination upon phylogenic reconstruction. Maximum likelihood (ML) trees were inferred using IQ-TREE v2.2.0.3, with 1000 ultrafast bootstrap (48, 49) under the GTR+F+I evolutionary model (empirical estimation of frequencies and allowing invariant sites). To improve within lineage relatedness inference, phylogenomic analyses were repeated on individual subsets of *R. salmoninarum* clades when relevant (i.e. 2022-24 BKD epizootic-related clusters). Visualization of trees and figures was done using the R software environment (R version 4.3.2) using the treedataverse 0.0.1. and ggplot2 packages.

### Data Deposition

## RESULTS

### Genome assemblies

Genome assemblies were constructed from short-read data from the 92 *R. salmoninarum* isolates sequenced in this study, along with 109 publicly available short-read datasets. The 201 genome assemblies generated contained 80-126 (average, 90) contigs, with N50 values ranging from 35,959 to 74,253 kbp (average, 64,074 kbp). Assembly sizes range from 2,981,688 to 3,078,808 kbp (average, 3,042,473 kbp), and GC content is 56.23-56.31% (average, 56.26%). Further quality metrics information for all the 201 genome assemblies is detailed in Table S2.

### Phylogenic reconstruction reveals an undescribed lineage of *R. salmoninarum* in the North-East Atlantic

The ANI values were above 99.75% for all pairs of isolates, demonstrating high genomic similarity. Phylogenomic analysis recovered three main lineages which were well supported with bootstrap values above 95% (UFBoot) (Fig. 2). An ML phylogeny was inferred based on a core genome alignment of 201 genomes comprising 2,788,055 nucleotide positions with an average nucleotide identity of 99.81% where the average genome coverage was 90.80%, as calculated by ParSNP. A total of 2,901 core-genome SNPs were identified between the most dissimilar *R. salmoninarum* isolates analyzed. The intra-lineage SNP distribution varied among the three lineages, with Lineage 1 exhibiting 395 SNPs between its most distant isolates, while Lineages 2 and 3 had 51 and 34 SNP differences, respectively. Inter-lineage comparisons revealed a minimum of 2,072 SNP sites between Linage 1 and Lineage 2, 2,822 SNP sites between Linage 1 and Lineage 3, and 2,389 SNP sites between Linage 2 and Lineage 3 (Fig. S2).

**Fig 2.**
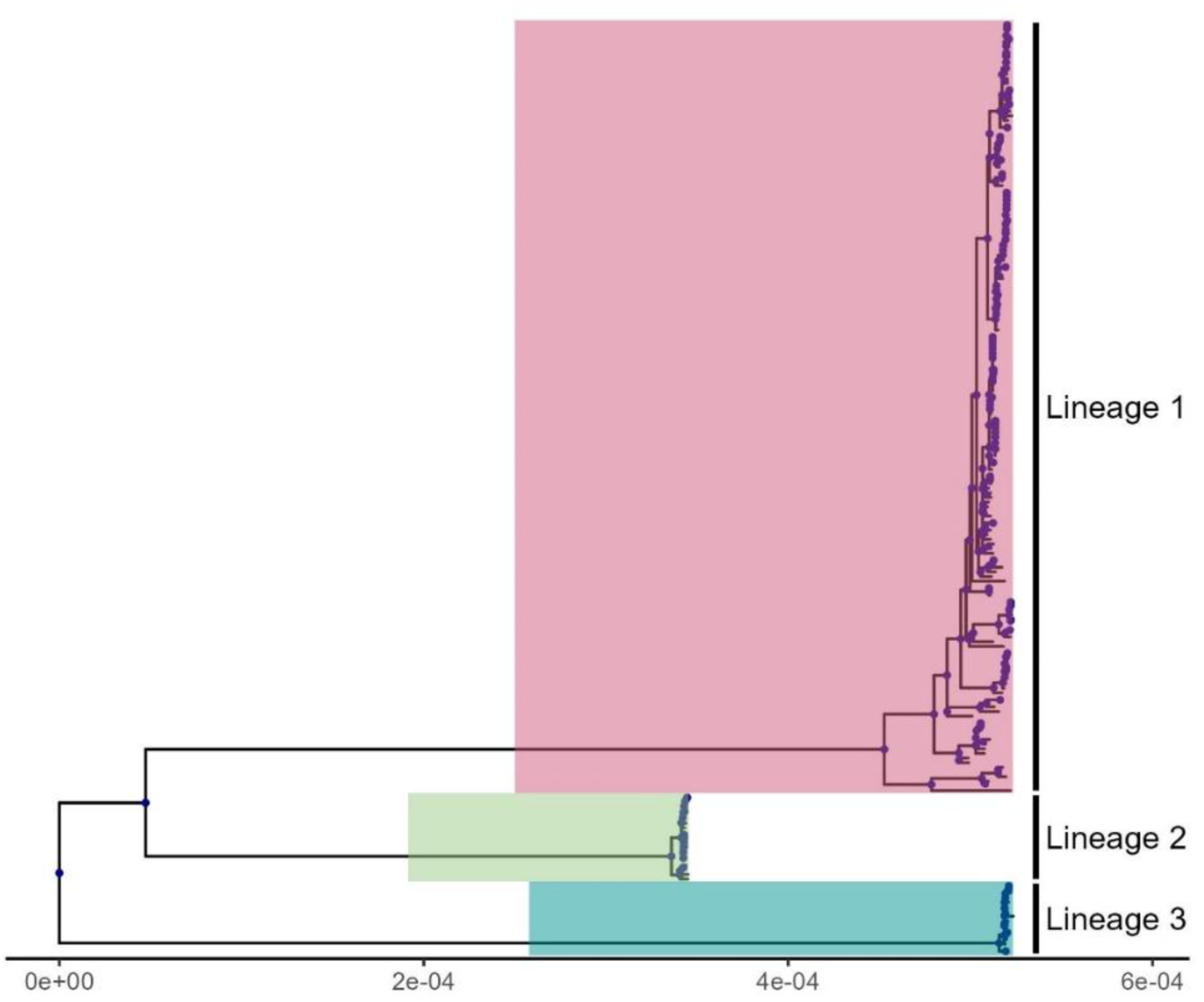
Maximum Likelihood (ML) core genome phylogenomic reconstruction of a global collection of 201 genomes showing the three main lineages of *Renibacterium salmoninarum*. Branch lengths are proportional to the number of nucleotide substitutions per site, as indicated by the scale bar at the bottom. The tree was midpoint rooted for visualization purposes. Dots at nodes represent bootstrap values ≥95.

Lineage 1 included 166 out of the 201 isolates, spanning more than 60 years of sampling (1960s to 2024). Isolates from all countries and host species were represented in Lineage 1 except for Iceland and the Faroe Islands, and Artic char, respectively (Fig. 3A). All 12 Danish and 47 of the Norwegian isolates sequenced were placed in Lineage 1. Among them, we identified three clades with high bootstrap support (UFBoot ≥95%) restricted to Norway: 1) Clade N1, with 16 isolates sampled between 2016 and 2024 in PA4 and PA5; 2) Clade N2, with 30 isolates sampled between 1997 and 2024 in PA6 and PA9 (single isolate); and 3) clade N3, with eight isolates sampled between 1998 to 2019 in PA3 and PA4. Moreover, nine of the Danish isolates formed Clade D1 (UFBoot ≥95%) sampled between 2017 and 2022 from different RAS facilities in Jutland, Denmark. These clades are annotated in Fig. 3A. Lineage 2 included the two original isolates from 1962 recovered from wild fish in the River Dee, Aberdeen. The remaining 17 isolates originated from Norway, with host species limited to the genus Salmo. The Norwegian isolates were geographically spread along the Norwegian coastline within production areas 3, 4, 5, 6, and 7; most were collected between 1985 and 1987, with one case in 1998 and the latest case in 2012 (Fig. 3B). In addition to Lineages 1 and 2, a third lineage of *R. salmoninarum* (Lineage 3, see Fig. 2) was identified in this study. This lineage consisted of sixteen isolates newly sequenced from the North-East Atlantic region, including all twelve *R. salmoninarum* from Iceland, one from the Faroe Islands, and three from Norway. Most Icelandic isolates were sampled between 1987 and 1990, with one isolate in 1996 and the latest in 2012, from Atlantic salmon, Arctic char, and rainbow trout. The sole *R. salmoninarum* isolate from the Faroe Islands was collected from Atlantic salmon before 1992. The Norwegian isolates were collected from Atlantic salmon in PA3 in 2005, and in PA13 in 2009 and 2012 (Fig. 3C).

**Fig 3.**
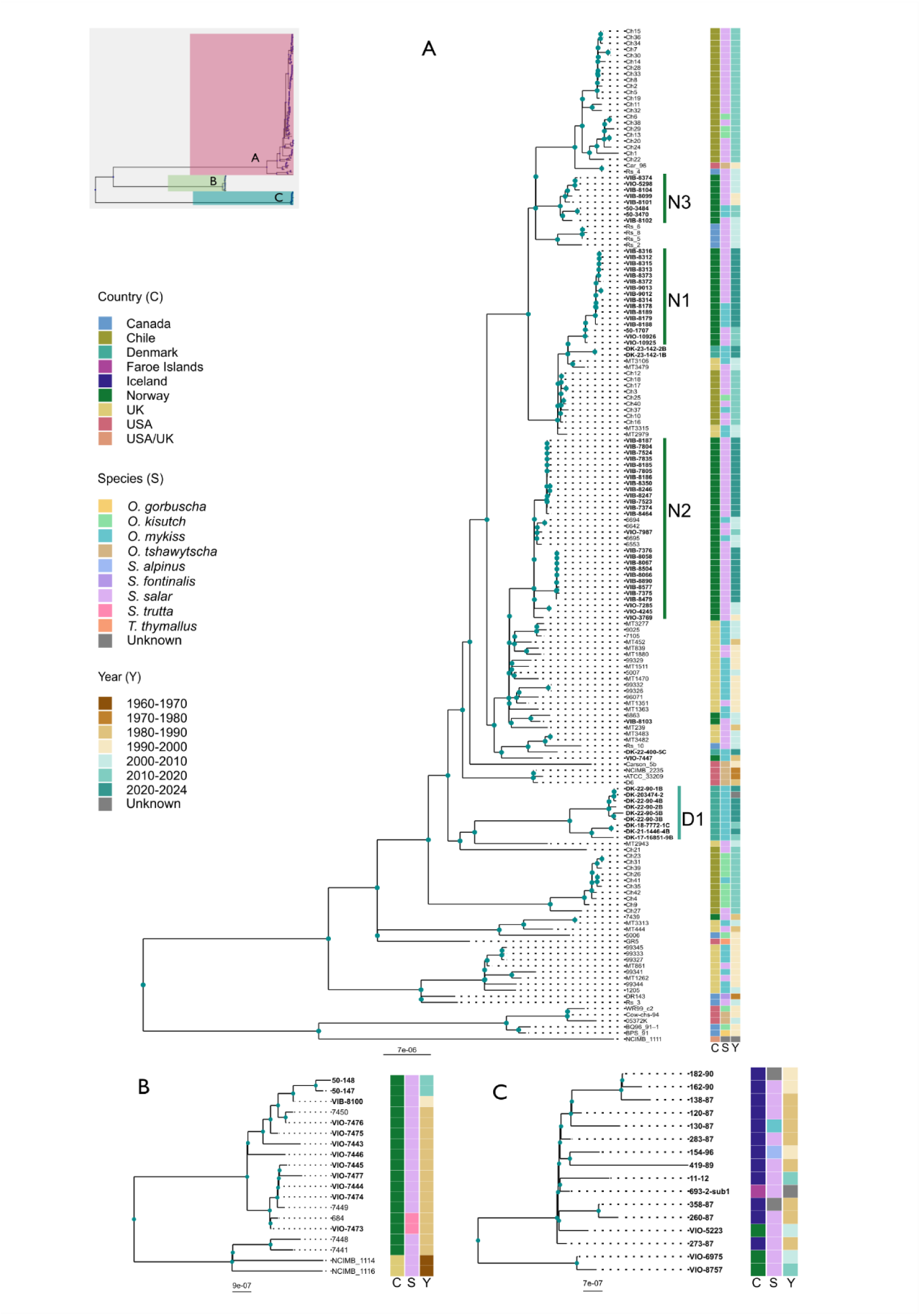
The top-left panel shows the Maximum Likelihood (ML) tree inferred from the core genome of a global collection of 201 genomes, with the three main lineages of *Renibacterium salmoninarum* colored. For clarity, each lineage is shown in panels A (Lineage 1), B (Lineage 2), and C (Lineage 3). Bootstrap support values at nodes in each of the three subtrees were 100%. Isolates sequenced in this study are shown in bold, and clades N1, N2, N3 (Norway), and D1 (Denmark) are annotated. Colored columns indicate associated metadata for country of origin (C), host species (S), and year of isolation (Y). Branch lengths are proportional to the number of nucleotide substitutions per site, as indicated by the scale bar at the bottom of the trees. Dots at nodes represent bootstrap values ≥95. Note: The previously published sequence for 5223 (ERR327964) was re-sequenced in our study. Phylogenomic analysis placed it at a distant position compared to the original sequence. To confirm this discrepancy, we performed additional re-sequencing, which confirmed its distant placement; therefore, the original public sequence was discarded and only our re-sequenced isolate was used.

### Two independent *R. salmoninarum* clades drive the current BKD outbreaks in Norway

*R. salmoninarum* has been present in Norway for more than four decades. Our results showed that the three above-described lineages have contributed to BKD cases along the Norwegian coastline. At least 7 of the 13 production areas in the country have registered detections of *R. salmoninarum*, with several of the cases yielding associated isolates dating back to the 1980s that we have analyzed herein (Fig. 1). Of the 67 isolates sequenced from Norway in the present study, 35 originated from farms within PA4 (n=11), PA5 (n=2), and PA6 (n=22) between December 2022 and August 2024. The geographical origins of the isolates involved are mapped in Fig. 4. These isolates resolved into two independently evolving clades within Lineage 1, both strongly supported (UFBoot ≥95%) and separated by a minimum pairwise distance of 66 SNPs. To enhance the resolution, core genome phylogenic analysis was applied separately to each clade.

**Fig 4.**
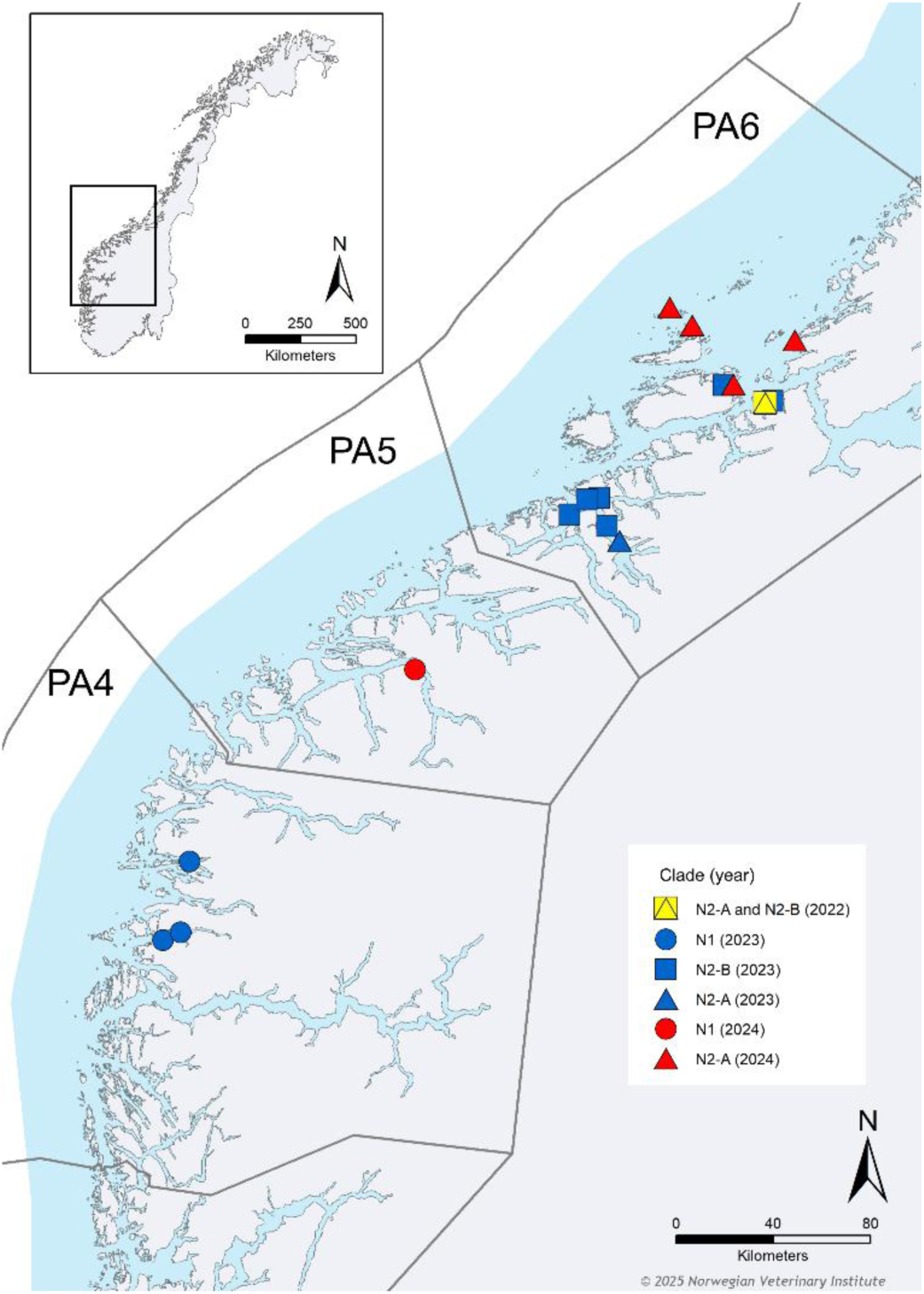
Map showing the location, year, and phylogenic division within Norwegian clades N1, N2-A, and N2-B of *Renibacterium salmoninarum*. The clades reflect distinct phylogenetic subdivisions within the BKD cases in PA4, PA5, and PA6 analyzed from 2022-2024. Clade N1 and Clade N2 were identified within the Maximum Likelihood (ML) tree inferred from the core genome of a global collection of 201 genome sequences. ML core genome phylogenomic analysis was performed independently for Clade N1 and Clade N2 subsets of genome sequences, which identified two strongly supported subclades N2-A and N2-B (UFBoot ≥95%). Modified from Attila Tarpai, the Norwegian Veterinary Institute. PA=Production area.

#### Clade N1

Clade N1 comprises 16 isolates from Atlantic salmon farms in PA4 (n=14), sites PA4-A to E, and PA5 (n=2) with one affected site, PA5-A. From PA4, three isolates were collected during 2016-2017 from sites PA4-A and PA4-B (two cases), and 11 recent isolates (2023) from three cases in sites PA4C-D; PA5 isolates originated from a case in PA5-A in 2024. Phylogenetic clustering generally corresponds to sampling location. The only apparent exception is VIB-8314 from PA4-E; however, the split between the isolate and other isolates from the same site had a low bootstrap value and was thus not supported by the phylogenomic tree (Fig. 5A-B).

**Fig 5.**
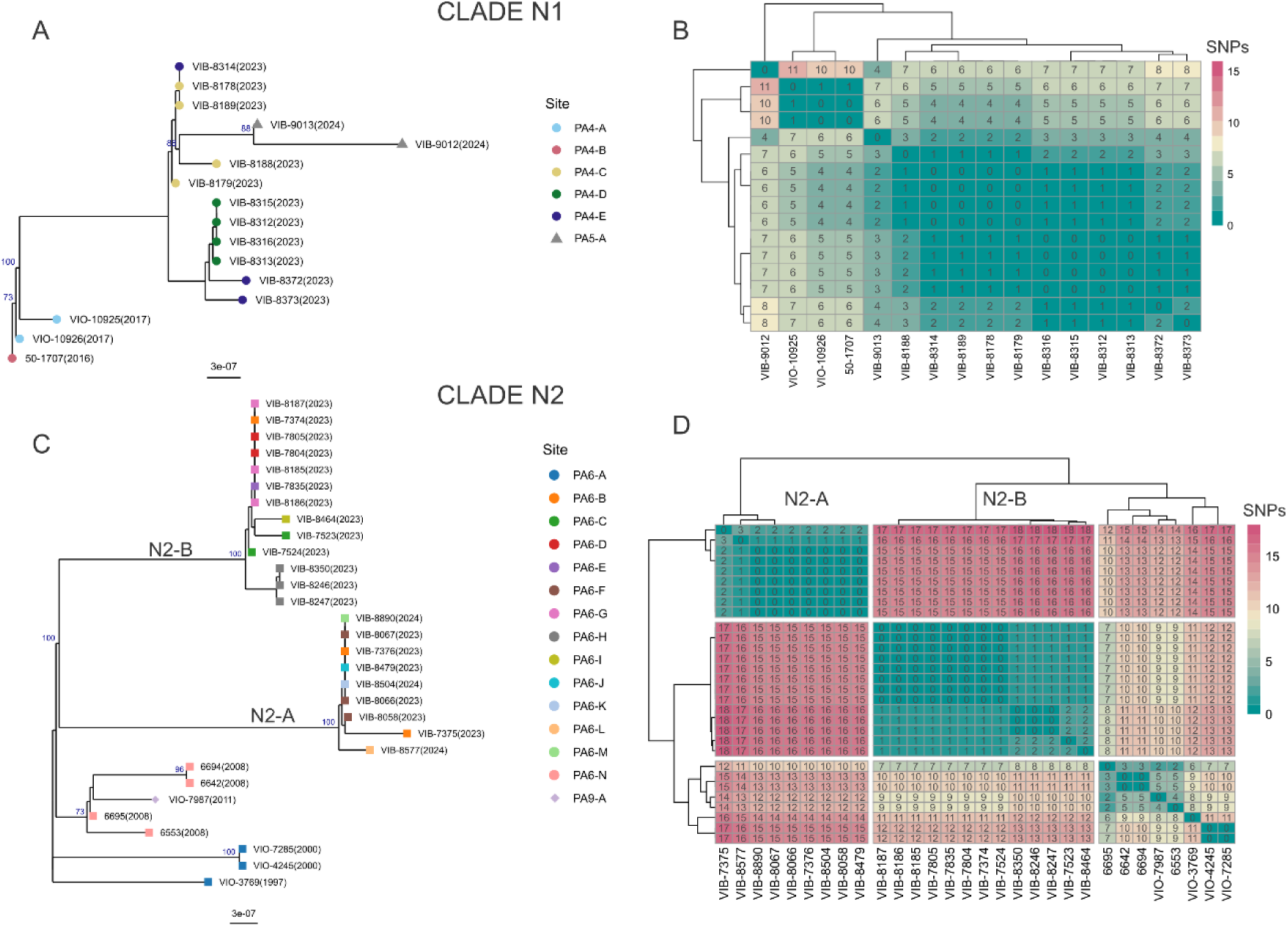
Maximum Likelihood (ML) core genome phylogenomic analysis was performed independently for Clade N1 (A) and Clade N2 (C) subsets of genome sequences. The trees were manually re-rooted in R using ggtree, setting the root at the node separating the oldest group of isolates within each clade for visualization purposes. Bootstrap values ≥70 are displayed at each node. Colored dots at the tips of the trees represent different sampling sites, and shapes indicate different production areas (PA). Collection year is shown in parentheses. Heatmaps illustrate SNP distance of the inferred core genomes, shown in each box for Clade N1 (B) and Clade N2 (D). Subclades N2-A and N2-B are annotated in the subtree (C) and heatmap (D) for Clade N2.

#### Clade N2

Clade N2 comprises 30 isolates, including 22 recent (2022–2024) and eight historical (1997–2011) isolates, predominantly from PA6 (n=29), sites PA6-A to N, with a single isolate from PA9 (n=1). Recent isolates represent 15 cases in Atlantic salmon farming sites across PA6: four in southwestern regions (PA6-D, PA6-E, PA6-F, and PA6-H) and eight in central areas (PA6-B, PA6-C, PA6-G, PA6-I, PA6-J, PA6-K, PA6-L, PA6-M). Phylogenomic analysis resolved two strongly supported subclades (N2-A: UFBoot 95%, N2-B: UFBoot 97%) separated by 15 to 18 core genome SNP differences (Fig. 5C-D). Even though both subclades exhibited short internal branches (N2-A: ≤3 SNP differences; N2-B: ≤2 SNP differences), they were not consistent with the geographical origins of the isolates (see Fig. 4). For example, three 2023 isolates collected on the same day from PA6-B (central PA6) were found in both subclades: the isolates VIB-7375 and VIB-7376 (N2-A) differed from VIB-7374 (N2-B) by 17–15 SNPs. In contrast, N2-B isolates VIB-7374 (PA6-B, central PA6) and VIB-7805 (PA6-D, southwestern PA6) were identical despite spatial separation. Within this clade we also found historical isolates which included those originating from PA6-N in 2008, and three from wild Atlantic salmon caught in the same area in the Orkla river (PA6-A; 1997, 2000). The 2000 Orkla isolates (VIO-4245 and VIO-7285) were identical but differed by 11 core genome SNPs from the 1997 isolate (VIO-3769). A single PA9 isolate from 2011 (VIO-7987, PA9-A), towards the north of the country, differed by 2-4 SNPs from the PA6-N isolates from 2008.

## DISCUSSION

The main objective of this study was to examine the epidemiological links of recent BKD cases using phylogeny-based analysis of genomic relatedness and to explore potential transmission patterns within affected areas. We posit that the current epizootics in western and mid regions of Norway, ongoing since 2022, are part of two independent transmission chains originating within Lineage 1. This work also reports the first *R. salmoninarum* genome sequences from Denmark, which likewise belonged to Lineage 1. Furthermore, our analysis identified a novel Lineage 3, comprising isolates predominantly from Iceland, the first publicly available genome from the Faroe Islands, and three isolates from Norway.

Whole genome-based analyses have been widely used for infectious disease outbreak assessments and are able to detect microevolution within clinically relevant, slow evolving microorganisms such as *Mycobacterium tuberculosis* (50). In accordance with the commonly used threshold of ANI levels of 95-96% for bacterial species demarcation (51), *R. salmoninarum* is the sole species described to date within the genus *Renibacterium*. The high pairwise ANI values (≥99.75%) among the isolates in this study, despite covering large spatiotemporal spans in origin, support the established understanding of the clonal population structure and the highly conserved genome of *R. salmoninarum* (36), which necessitate use of high-resolution methods such as WGS to be able to differentiate between isolates.

### A novel genetic pool of *R. salmoninarum* in the North-East Atlantic

Our phylogenomic analysis confirms the previously described division of *R. salmoninarum* into two major lineages and further identifies a third, previously unrecognized lineage. Global spread facilitated by growing aquaculture-related activities has been suggested for other aquatic pathogens such as *Yersinia ruckeri*, whose population structure shows an intricate interplay between historic range expansion and more recent migration due to human activities (52, 53). Anthropogenic aquaculture operations were established as a major driver for the intercontinental spread and expansion of *R. salmoninarum* Lineage 1 (26, 37). Indeed, in our study, Lineage 1 exhibited notably greater genomic heterogeneity than Lineages 2 and 3 (Fig. S2), although this may simply reflect the over-representation of aquaculture-associated BKD cases in the sampling. Future studies incorporating wild fish isolates would help disentangle true lineage diversity from surveillance artifacts. In contrast to the widespread distribution of Lineage 1, Lineage 3 is composed almost exclusively by Icelandic isolates and does not appear to have spread beyond the North-East Atlantic region.

### Two transmission chains driving current BKD epizootics in Norway

BKD had only been sporadically (0-3 cases per year) reported in Norway for several years, before cases increased markedly in late 2022 (Fig. 1). The initial cases prompted increased sampling in the affected production areas in an attempt to understand and control the spread of the pathogen. The first reported cases emerged in PA6-B (December 2022), followed by detection at the neighboring PA6-C site (February 2023). Subsequent cases occurred in PA6 and PA4, with the most recent cases in this study detected in PA5 (PA5-A) during the summer of 2024. All recent Norwegian isolates belonged to Lineage 1, forming two distinct clades and exhibiting strong spatial confinement: A) Clade N1 (cases from PA4 and PA5) and B) Clade N2 (cases from PA6), see Fig. 4 and 5. These findings showcase two genomically distinct groups of isolates, suggesting two independent emergences followed by local transmission. The exact route by which the bacteria reached the fish in our study remains unclear, and several introductions to the sites in PA4 and PA5 rather than direct transmission between them cannot be excluded. Epidemiological data involving the transport of fish across production areas where BKD was recently detected are consistent with the possible role of sea site operations in the spread of the disease (54). Within PAs, we have documented genomically diverse variants coexisting at an individual site, as well as different sites sharing identical isolates, as illustrated in Fig. 4. Similar patterns were reported by Bayliss et al. in Chile, which described both persistence of highly similar isolates and multi-variant colonization at particular sites (37). Phylogenomic relationships among isolates was not reflected in or associated with company affiliation (data not shown), consistent with findings from Chilean studies where pathogen transmission occurred through the aquaculture network regardless of the company and the production stage (37).

### Historical transmission and endemicity in the North-East Atlantic waters

Our study suggests that *R. salmoninarum* movements across production areas occurred in the past, and that the bacterium persisted and sometimes re-emerged in specific regions after years of apparent absence, challenging assumptions of pathogen elimination and underscoring the risk of resurgence. An example of historical potential transmission across distant production areas is provided by Clade N2, which contains isolates almost exclusively from PA6, except for a single isolate from PA9 (2011) highly related to isolates from a case in PA6 in 2008 (Fig. 5C-D). Likewise, Clade N3 (see Fig. 3A) illustrates possible transmissions between PA3 and PA4, in this case over a span of two decades. This clade includes cases registered in 2003 and 2005 that were not detected in the region thereafter, and which are phylogenomically distinct from more recent cases in PA4 (Clade N1). Interestingly, genomically similar isolates within Clade N3 were identified in the adjacent PA3 as early as 1998, with subsequent occurrences in 2005 and again in 2019, suggesting persistence and potential re-emergence. These patterns likely reflect anthropogenic activities, such as the movement of fish or ova, a mechanism similarly implicated in the pathogen spread observed within Chilean aquaculture networks (37), or low-level circulation within wild fish in the areas. Furthermore, Clade N1 illustrates the persistence or reintroduction of closely related strains, as contemporary isolates from PA4 and PA5 are highly similar to those collected during 2016 and 2017 in PA4 from farmed salmon (see Fig. 5A). In addition, our study reveals close genomic relationships between wild fish isolates in Clade N2, suggesting that *R. salmoninarum* may have persisted in wild Atlantic salmon populations in the Orkla river, PA6-A, at least between 1997 and 2000 (see Fig. 5C). These wild fish isolates were closely related to those from recent cases collected from farmed Atlantic salmon in the same region. Subclinical carriage of *R. salmoninarum* within fish populations has been well documented (11, 17, 55, 56), and Johansen et al., 2011, highlighted the risk for transmission between wild and farmed populations (57). Altogether, these findings are consistent with the existence of endemic reservoirs of *R. salmoninarum* which would facilitate the persistence and local spread of the bacterium (16, 17, 26, 37). The sudden increase in BKD cases in Norway underscores the need to investigate stressors that may trigger proliferation of the pathogen if present at subclinical levels. This emphasizes the need for continuous monitoring to control disease.

*R. salmoninarum* is likely endemic in any country with native salmonid populations or active salmonid aquaculture, and BKD has been described as a long-term endemic disease in European waters (26, 56). Lineage 2 remains restricted to isolates from Scotland and Norway, with early cases in Norway being attributed to transmission from wild populations of the North Sea (26). Notably, the limited number of recent Norwegian isolates recovered during the years following the implementation of biosecurity measures in the 1990s (18, 26, 57) appear to have efficiently curbed further spread of the lineage. Indeed, none of the subsequent detections in Norway belonged to Lineage 2.

The discovery of Lineage 3, encompassing all Icelandic isolates, represents a significant finding. *R. salmoninarum* is frequently reported in Icelandic wild fish through routine surveillance and is regarded as endemic to the country (17). The first available *R. salmoninarum* genome sequence from the Faroe Islands also belongs to this lineage. This could indicate either circulation of *R. salmoninarum* from the same genetic pool present in Icelandic waters, or its introduction through commercial activities in the past. The latter hypothesis is likely to explain the presence of this lineage in Norway, which is represented by three isolated cases. The 2012 Norwegian Fish Health Report suggested that a potential introduction may have occurred through salmonid import activities in 2011 (58). Like most countries, Iceland has relied on diagnostic methods for BKD other than bacterial culture since the early 1990s, due to the challenges in cultivating *R. salmoninarum*. The lack of recent Icelandic and Faroese isolates in our collection limits our ability to explore the Lineage 3 in greater detail. However, given the genomic distinctiveness of these isolates compared to those from Lineages 1 and 2, it seems reasonable to propose that a natural reservoir of Lineage 3 has existed for a long time in the North-East Atlantic and might still remain. Our findings underscore the well-known host switching ability of *R. salmoninarum*, consistent with those of Brynildsrud et al. for Lineage 1 (26). This is illustrated by Lineage 3 isolates from Iceland, which exhibit minimal genetic variation yet were collected from three different Icelandic salmonid species.

### Persistence of *R. salmoninarum* within natural reservoirs

The emergence of infectious diseases in aquaculture often stems from unclear drivers as discussed by Murray & Peeler (59). The extent to which such emergence is shaped by either strong selection in farming systems and/or ‘opportunistic’ infections from environmental reservoirs, varies based on ecology, disease dynamics, and pathogen etiology (35). While in the present study the spread of *R. salmoninarum* within and across the affected areas can likely be linked to commercial activities, the chronic nature of BKD (60, 61) and its typical subclinical presentation (17, 62) are important factors to consider when exploring the potential source(s) of BKD cases. The ability of *R. salmoninarum* to persist in different environments has to some extent been investigated. It has been shown to survive in sediment and fecal material for up to 21 days (63), and recent Swedish field studies (56) additionally highlight its potential for environmental persistence, particularly in areas with open aquaculture systems. Here, the presence of older, and genomically similar isolates to those driving the recent epizootics within the N1 and N2 clades, supports the hypothesis that natural reservoirs and/or ongoing, low-level circulation of the pathogen exist within specific regions. This finding raises the possibility of epidemiological continuity between historical and recent cases, yet the existence and nature of such a link cannot be ascertained from the present data.

Understanding the implications intracellular lifestyles may have on environmental persistence is key to inform water treatment and disinfection procedures to more effectively interrupt the horizontal and vertical transmission cycles in aquaculture facilities. The intracellular nature of *R. salmoninarum* represents a complex evolutionary adaptation that may also enhance its environmental persistence. Other aquaculture intracellular pathogens have shown extended extracellular viability likely stemming from adaptations originally evolved to survive within the host cells (64–66). Characteristic traits of *R. salmoninarum* such as its durable cell wall (61) protecting against degradation within macrophages (67), its potential ability to withstand nutrient limitations within the host cells (14), and its slow growth rate (14) might facilitate survival in the external environment. It has been proposed that *R. salmoninarum* may enter a dormant state when exposed to suboptimal conditions (14, 61, 68) which could be an effective survival strategy until reaching a new host. Moreover, the intracellular lifestyle of this bacterium contributes to its ability for vertical transmission (8), which has broad implications for aquaculture biosecurity. In this context, broodstock screening and management become critical control points, as infected mature fish can transmit the pathogen to multiple generations (60).

### Aquaculture practices and biosecurity challenges

Commercial aquaculture, with the large-scale trade and transport of fish, eggs, and feed across wide geographical regions may facilitate the transmission of pathogens (2), as seen in the intercontinental transmission of *R. salmoninarum* (26, 37). A critical stage in salmon production is the transport of smolts by well-boats, which should adhere to an “all in, all out” principle, including disinfection between loads and strict separation from boats used for transport to slaughter. Current sea site operations in Norway involve significant interaction among different regions, companies, and sites due to the shared use of hatcheries, well-boats, delousing vessels, and service boats (54). Notably, operations across multiple sites in PA6 coincided with BKD spread in our study, despite certified disinfection protocols (54). Washing and disinfection help reduce the risk of pathogen dissemination but cannot eliminate it entirely, and limited access to well-boat systems may also come in conflict with time-consuming thorough cleaning regimens (69). While the risk of infection from pathogen exposure during transport is not definitely established, the potential for well-boats contaminated with pathogens to act as vectors for environmental transmission should be considered (70).

In the case of Denmark, several outbreaks of BKD have been registered (71) since the first detection of *R. salmoninarum* in the Skjern Å river system in the late 1990s (20), and this study provides the first full genome sequences from Danish isolates. These isolates were recovered from recent BKD outbreaks in rainbow trout mainly from land-based RAS farms and belong exclusively to the aquaculture-related Lineage 1. BKD is perceived as an important disease in RAS systems in Denmark. Once the infection enters a facility, successful disinfection can be challenging due to the capability of the bacterium to persist in the complex piping systems of such farms (72). While strict biosecurity measures should maintain a disease-free status, pathogens can enter through contaminated water, equipment, or personnel contact if any of these measures fail or are not properly implemented, both in well-boats (2, 70) and in RAS (73, 74). In a study on risk factors in RAS and well-boat systems, insufficient knowledge was reported as a common contributor to biosecurity breaches (70). Our results underscore the importance of revising and optimizing disinfection protocols for *R. salmoninarum* to reduce the risk of future outbreaks.

The scope and interpretation of our study should consider some limitations intrinsic to the collection of *R. salmoninarum* isolates, including the non-representative sampling across Norwegian production areas and a lack of isolates from some of the other countries with significant salmonid aquaculture industries. While our targeted sampling approach enhanced outbreak study accuracy, broader geographical gaps in sampling and the reliance on identification methods other than bacterial culturing in some countries (such as the cases from Denmark and Iceland) limited our ability to generate a more comprehensive whole-genome sequence data set. Bacterial culture on selective media is extremely slow, which can delay BKD diagnosis and can fail to detect BKD especially from subclinical infections (14, 62). This slow and fastidious growth of *R. salmoninarum* makes the acquisition of pure cultures for WGS particularly challenging, presenting a significant obstacle for genomic studies. This highlights the need for alternative approaches that do not rely solely on culture-based methods for the detection of *R. salmoninarum*. A consistent surveillance based on efficient and standardized BKD detection methods is crucial for us to better understand *R. salmoninarum* distribution and evolution patterns.

## CONCLUSION

The discovery of a previously unrecognized, genetically distinct lineage of *R. salmoninarum*, almost exclusively Icelandic, represents a significant advance in our understanding of the evolutionary history of the pathogen and its geographic distribution across the North-East Atlantic. WGS enabled the identification of two distinct *R. salmoninarum* clades underlying the recent epizootics in Norway, supporting the hypothesis that multiple endemic reservoirs of *R. salmoninarum* exist within European water systems. Specific strains appear to have established persistent populations in defined geographic regions following their introduction, with indications for transmission between wild and farmed populations and bacterial persistence in aquatic environments, likely via asymptomatic carrier fish. We propose plausible pathways of local spread within Norway, primarily associated with anthropogenic activities such as fish transport and well-boat operations, potentially facilitated by inadequate biosecurity procedures. The sudden emergence of BKD in Norway without a clear source of introduction highlights the need to investigate both the mechanisms by which *R. salmoninarum* persist at subclinical levels, and the factors that can trigger its activation. Ongoing surveillance and targeted management strategies will be critical to mitigate future outbreaks and protect both wild and farmed salmonid populations.

## ACKNOWLEDGMENTS

This work was co-funded by the European Union’s Horizon Europe Project 101136346 EUPAHW, in addition to A. Villamil-Alonso Ph.D. research project granted by Technical University of Denmark, DTU n° 102184, the National Reference Laboratory for Fish and Crustacean Diseases of Denmark (NRL), and the Henriksen’s fund grant 2021-05-24-153856 for improved diagnostic of Bacterial Kidney Disease (BKD) in rainbow trout.

We would like to thank Eve Marie Louise Zeyl Fiskebeck for her critical revision of the paper.

---

**Supplementary Figure 1.**
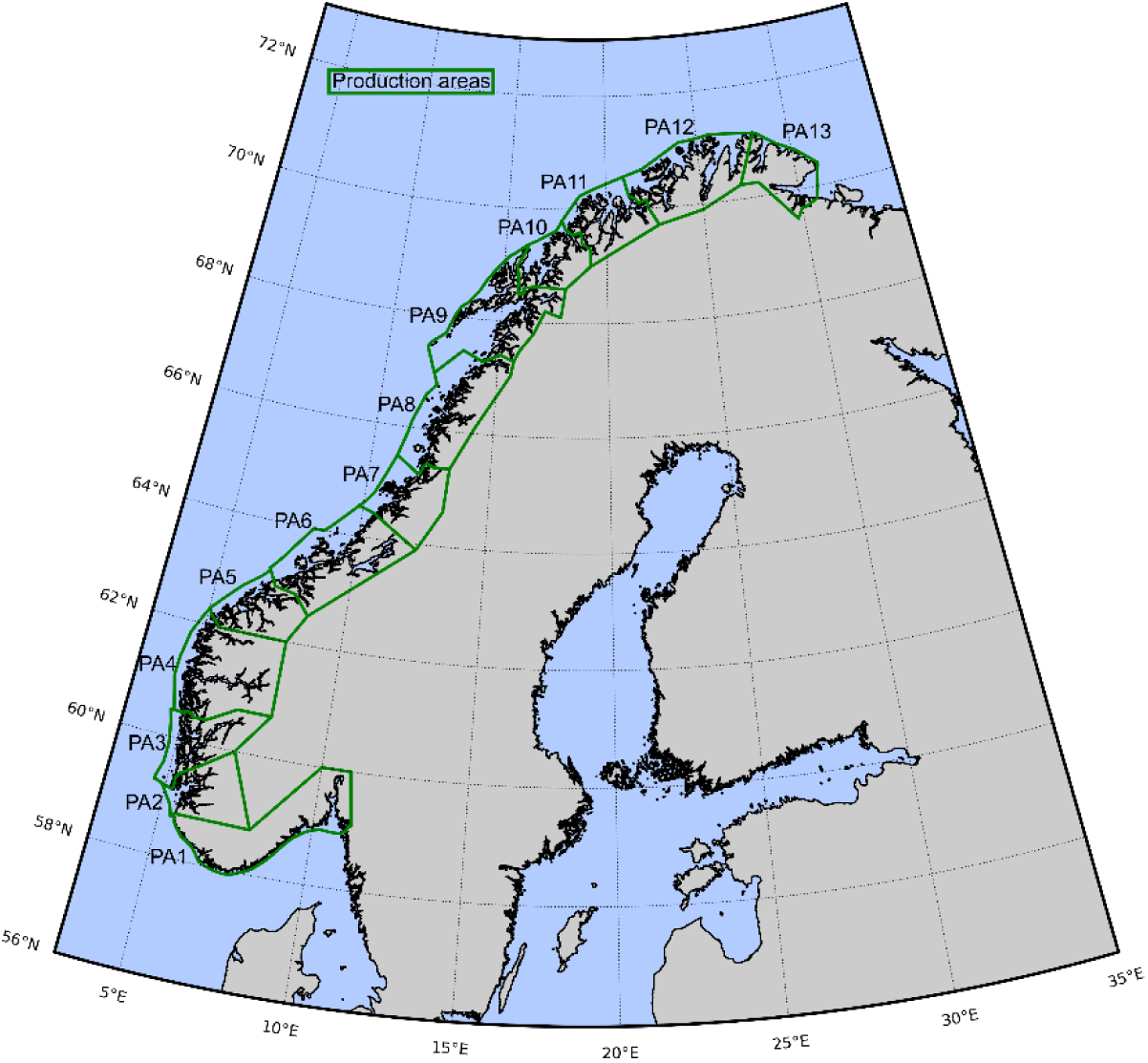
Production zones 1-13 along the Norwegian coast modified from Lovdata, 2017.

**Supplementary Figure 2.**
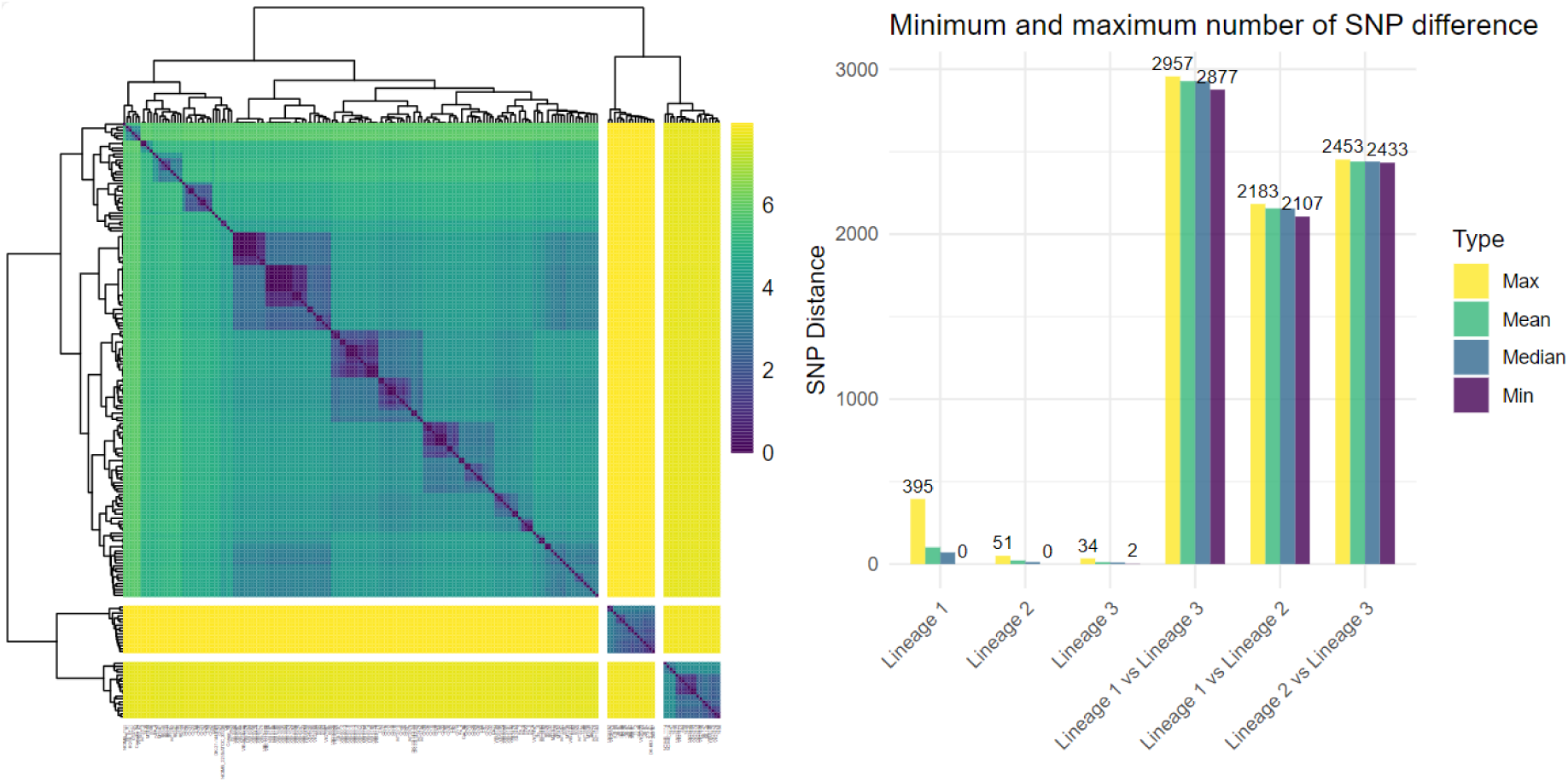
Left: Heatmap displaying the single nucleotide polymorphism (SNP) distance matrix for a global collection of 201 *Renibacterium salmoninarum* genomes. Values were log-transformed, and hierarchical clustering was applied for visualization purposes. Right: Minimum and maximum SNP differences. Yellow, green, blue, and purple bars represent the maximum, median, mean, and minimum SNP differences between isolates within lineages and between lineages.

## REFERENCES

1. Food and Agriculture Organization of the United Nations. 2024. The state of world fisheries and aquaculture 2024. Food and Agriculture Organization of the United Nations, Rome, Italy. https://openknowledge.fao.org/handle/20.500.14283/cd0683en.

2. Naylor RL, Hardy RW, Buschmann AH, Bush SR, Cao L, Klinger DH, Little DC, Lubchenco J, Shumway SE, Troell M. 2021. A 20-year retrospective review of global aquaculture. Nature 591:551–563.

3. Fryer JL, Sanders JE. 1981. Bacterial kidney disease of salmonid fish. Annu Rev Microbiol 35:273–298.

4. Banner CR, Rohovec JS, Fryer JL. 1991. A new value for mol percent guanine + cytosine of DNA for the salmonid fish pathogen *Renibacterium salmoninarum*. FEMS Microbiol Lett 63:57–59.

5. EFSA Panel on Animal Health and Welfare (AHAW), Nielsen SS, Alvarez J, Calistri P, Canali E, Drewe JA, Garin-Bastuji B, Gonzales Rojas JL, Gortázar C, Herskin MS, Michel V, Miranda Chueca MÁ, Padalino B, Roberts HC, Spoolder H, Ståhl K, Velarde A, Viltrop A, Winckler C, Bron J, Olesen NJ, Sindre H, Stone D, Vendramin N, Antoniou SE, Aznar I, Papanikolaou A, Karagianni AE, Bicout DJ. 2023. Assessment of listing and categorisation of animal diseases within the framework of the Animal Health Law (Regulation (EU) 2016/429): Bacterial kidney disease (BKD). EFSA J 21:e08326.

6. Balfry S, Albright L, Evelyn T. 1996. Horizontal transfer of *Renibacterium salmoninarum* among farmed salmonids via the fecal-oral route. Dis Aquat Organ 25:63–69.

7. Evenden AJ, Grayson TH, Gilpin ML, Munn CB. 1993. *Renibacterium salmoninarum* and bacterial kidney disease — the unfinished jigsaw. Annu Rev Fish Dis 3:87–104.

8. Evelyn TPT, Ketcheson JE, Prosperi-Porta L. 1984. Further evidence for the presence of *Renibacterium salmoninarum* in salmonid eggs and for the failure of povidone-iodine to reduce the intra-ovum infection rate in water-hardened eggs. J Fish Dis 7:173–182.

9. Pascho RJ, Elliott DG, Streufert JM. 1991. Brood stock segregation of spring chinook salmon *Oncorhynchus tshawytscha* by use of the enzyme linked immunosorbent assay (ELISA)and the fluorescent antibody technique (FAT) affects the prevalence and levels of *Renibacterium salmoninarum* infection in progeny. Dis Aquat Organ 12:25–40.

10. Riepe TB, Fetherman ER, Neuschwanger B, Davis T, Perkins A, Winkelman DL. 2023. Vertical transmission of *Renibacterium salmoninarum* in cutthroat trout (*Oncorhynchus clarkii*). J Fish Dis 46:309–319.

11. Murray AG, Munro LA, Wallace IS, Allan CET, Peeler EJ, Thrush MA. 2012. Epidemiology of *Renibacterium salmoninarum* in Scotland and the potential for compartmentalised management of salmon and trout farming areas. Aquaculture 324–325:1–13.

12. Banner CR, Long JJ, Fryer JL, Rohovec JS. 1986. Occurrence of salmonid fish infected with *Renibacterium salmoninarum* in the Pacific Ocean. J Fish Dis 9:273–275.

13. Hall LM, Duguid S, Wallace IS, Murray AG. 2015. Estimating the prevalence of *Renibacterium salmoninarum*-infected salmonid production sites. J Fish Dis 38:231–235.

14. Hirvelä-Koski V, Pohjanvirta T, Koski P, Sukura A. 2006. Atypical growth of *Renibacterium salmoninarum* in subclinical infections. J Fish Dis 29:21–29.

15. Bruno DW. 1986. Scottish experience with bacterial kidney disease in farmed salmonids between 1976 and 1985. Aquac Res 17:185–190.

16. Chambers E, Gardiner R, Peeler EJ. 2008. An investigation into the prevalence of *Renibacterium salmoninarum* in farmed rainbow trout, *Oncorhynchus mykiss* (Walbaum), and wild fish populations in selected river catchments in England and Wales between 1998 and 2000. J Fish Dis 31:89–96.

17. H. Jonsdottir, H.J. Malmquist, S.S. Snorrason, G. Gudbergsson, S. Gudmundsdottir. 1998. Epidemiology of *Renibacterium salmoninarum* in wild Arctic charr and brown trout in Iceland. J Fish Biol 53:322–339.

18. Binde M. 2023. Nytten av regelverk ved kontroll av BKD, p. 40–45. *In* Norsk Veterinærhistorisk Selskap Årbok 2023.

19. Håstein T, Dale OB. 1987. Registrations of bacterial kidney disease in Norway 1980–87, p. 8. *In* Proceedings of the Third International Conference. Bergen, Norway.

20. Lorenzen E, Olesen NJ, Korsholm H, Heuer OE, Evensen Ø. 1997. First demonstration of *Renibacterium salmoninarum* / BKD in Denmark. Bull Eur Assoc Fish Pathol 17:140–144.

21. Kristmundsson Á, Árnason F, Gudmundsdóttir S, Antonsson T. 2015. Levels of *Renibacterium salmoninarum* antigens in resident and anadromous salmonids in the River Ellidaár system in Iceland 10.1111/jfd.12401.

22. Rhodes L, Durkin C, Nance S, Rice C. 2006. Prevalence and analysis of *Renibacterium salmoninarum* infection among juvenile Chinook salmon *Oncorhynchus tshawytscha* in North Puget Sound. Dis Aquat Organ 71:179–190.

23. Bruno DW. 2015. Bacterial kidney disease. ICES. Identification Leaflets for Diseases and Parasites of Fish and Shellfish. Leaflet No. 21. 7 pp.

24. Forseth T, Sigurd E, Fiske P, Falkegård M, Garmo ØA, Garseth ÅH, Skoglund H, Solberg MF, Thorstad EB, Utne KR, Vøllestad A, Vollset KW, Wennevik V. 2024. Status of wild Atlantic salmon in Norway 2024. 19. Trondheim.

25. Bøe K, Bjøru B, Tangvold Bårdsen M, Nordtug Wist A, Wolla S, Sivertsen A. 2021. Opportunities and challenges related to sperm cryopreservation in Atlantic salmon gene banks. Conserv Sci Pract 3:e552.

26. Brynildsrud O, Feil EJ, Bohlin J, Castillo-Ramirez S, Colquhoun D, McCarthy U, Matejusova IM, Rhodes LD, Wiens GD, Verner-Jeffreys DW. 2014. Microevolution of *Renibacterium salmoninarum*: evidence for intercontinental dissemination associated with fish movements. ISME J 8:746–756.

27. Lovdata. 2022. Regulations relating to animal health (Animal Health Regulations) Chapter II. Supplementary national provisions for aquatic animals – listing and notification obligation. Norwegian Food Safety Authority, Oslo, Norway.

28. Lovdata. 2017. Regulations relating to production areas for aquaculture of edible fish in the sea of salmon, trout and rainbow trout (Production Area Regulations). Norwegian Food Safety Authority, Oslo, Norway.

29. Dale OB, Håstein T, Reitan LJ, Faller R, Endal T, Heum M, Steinum. 2008. The surveillance and control programme for bacterial kidney disease (BKD) in Norway. 149. Oslo, Norway.

30. Garseth Å, Furnesvik L, Hansen H, Haukland H, Kielland Ø, Strand D. 2025. Villfiskrapporten 2024. Veterinærinstituttets rapportserie. Utgitt av Veterinærinstituttet 2025.

31. Moldal T, Wiik-Nielsen J, Oliveira V, Svendsen J, Sommerset I. 2025. Norwegian Fish Health Report 2024. Norwegian Veterinary Institute Report series. Norwegian Veterinary Institute.

32. A. Kristmundsson. 2022. Chapter 25 - Bacterial kidney disease, in: F.S.B. Kibenge, Baldisserotto, B., Sie-Maen Chong, R. (Ed.), Aquaculture Pathophysiology, Academic Press, 2022.

33. S. Guðmundsdóttir, H. Sigurjónsdóttir, S. Matthíasdóttir, H. Jónsdóttir, B. Laxdal, E. Benediktsdóttir. 2000. Measures applied to control *Renibacterium salmoninarum* infection in Atlantic salmon: A retrospective study of two sea ranches in Iceland. Aquaculture 186 (3–4):193–203.

34. European Commission. 2021. Commission Implementing Decision (EU) 2021/260 of 11 February 2021 approving national measures designed to limit the impact of certain diseases of aquatic animals in accordance with Article 226(3) of Regulation (EU) 2016/429 of the European Parliament and of the Council and repealing Commission Decision 2010/221/EU.

35. Bayliss SC, Verner-Jeffreys DW, Bartie KL, Aanensen DM, Sheppard SK, Adams A, Feil EJ. 2017. The promise of whole genome pathogen sequencing for the molecular epidemiology of emerging aquaculture pathogens. Front Microbiol 8.

36. Wiens GD, Rockey DD, Wu Z, Chang J, Levy R, Crane S, Chen DS, Capri GR, Burnett JR, Sudheesh PS, Schipma MJ, Burd H, Bhattacharyya A, Rhodes LD, Kaul R, Strom MS. 2008. Genome sequence of the fish pathogen *Renibacterium salmoninarum* suggests reductive evolution away from an environmental *Arthrobacter* ancestor. J Bacteriol 190:6970–6982.

37. Bayliss SC, Verner-Jeffreys DW, Ryder D, Suarez R, Ramirez R, Romero J, Pascoe B, Sheppard SK, Godoy M, Feil EJ. 2018. Genomic epidemiology of the commercially important pathogen *Renibacterium salmoninarum* within the Chilean salmon industry. Microb Genomics 4:e000201.

38. Austin B, Embley TM, Goodfellow M. 1983. Selective isolation of *Renibacterium salmoninarum*. FEMS Microbiol Lett 17:111–114.

39. Kaspersen H. 2025. NorwegianVeterinaryInstitute/Assemblage: v0.2.1 (v0.2.1). Zenodo. 10.5281/zenodo.15647694.

40. Ewels P, Magnusson M, Lundin S, Käller M. 2016. MultiQC: summarize analysis results for multiple tools and samples in a single report. Bioinforma Oxf Engl 32:3047–3048.

41. Bittencourt S. 2010. FastQC: A quality control tool for high throughput sequence data – ScienceOpen. https://www.scienceopen.com/document?vid=de674375-ab83-4595-afa9-4c8aa9e4e736.

42. Wood DE, Lu J, Langmead B. 2019. Improved metagenomic analysis with Kraken 2. Genome Biol 20:257.

43. Hall M. 2022. Rasusa: Randomly subsample sequencing reads to a specified coverage. J Open Source Softw 7:3941.

44. Wick R, Judd L, Gorrie C, Holt K. 2017. Unicycler: Resolving bacterial genome assemblies from short and long sequencing reads. PLOS Comput Biol 13:e1005595.

45. Gurevich A, Saveliev V, Vyahhi N, Tesler G. 2013. QUAST: quality assessment tool for genome assemblies. Bioinforma Oxf Engl 29:1072–1075.

46. Kaspersen H, Fiskebeck EZ. 2022. ALPPACA - A tooL for Prokaryotic Phylogeny And Clustering Analysis. J Open Source Softw 7:4677.

47. Nicholas J. Croucher, Andrew J. Page, Thomas R. Connor, Aidan J. Delaney, Jacqueline A. Keane, Stephen D. Bentley, Julian Parkhill, Simon R. Harris. 2014. Rapid phylogenetic analysis of large samples of recombinant bacterial whole genome sequences using Gubbins | Nucleic Acids Research | Oxford Academic. Nucleic Acids Res 43:e15.

48. Minh B, Schmidt H, Chernomor O, Schrempf D, Woodhams M, Haeseler A, Lanfear R. 2020. IQ-TREE 2: New models and efficient methods for phylogenetic inference in the genomic era. Mol Biol Evol Oxf Acad 37:2461.

49. Hoang D, Chernomor O, Haeseler A, Minh B, Vinh L. 2017. UFBoot2: Improving the Ultrafast Bootstrap approximation. Mol Biol Evol Oxf Acad 35:518–522.

50. Walker TM, Ip CL, Harrell RH, Evans JT, Kapatai G, Dedicoat MJ, Eyre DW, Wilson DJ, Hawkey PM, Crook DW, Parkhill J, Harris D, Walker AS, Bowden R, Monk P, Smith EG, Peto TE. 2013. Whole-genome sequencing to delineate *Mycobacterium tuberculosis* outbreaks: a retrospective observational study. Lancet Infect Dis 13:137–146.

51. Kim M, Oh H-S, Park S-C, Chun J. 2014. Towards a taxonomic coherence between average nucleotide identity and 16S rRNA gene sequence similarity for species demarcation of prokaryotes. Int J Syst Evol Microbiol 64:346–351.

52. Bastardo A, Ravelo C, Romalde JL. 2015. Phylogeography of *Yersinia ruckeri* reveals effects of past evolutionary events on the current strain distribution and explains variations in the global transmission of enteric redmouth (ERM) disease. Front Microbiol 6:1198.

53. Gulla S, Barnes AC, Welch TJ, Romalde JL, Ryder D, Ormsby MJ, Carson J, Lagesen K, Verner-Jeffreys DW, Davies RL, Colquhoun DJ. 2018. Multilocus Variable-Number Tandem-Repeat Analysis of *Yersinia ruckeri* Confirms the Existence of Host Specificity, Geographic Endemism, and Anthropogenic Dissemination of Virulent Clones. Appl Environ Microbiol 84:e00730–18.

54. Sommerset, I., Wiik-Nielsen J, Moldal T, Oliveira VHS, Svendsen JC, Haukaas A, Brun E. 2024. Norwegian Fish Health Report 2023. Norwegian Veterinary Institute Report.

55. Delghandi MR, Menanteau-Ledouble S, Waldner K, El-Matbouli M. 2020. *Renibacterium salmoninarum* and *Mycobacterium* spp.: two bacterial pathogens present at low levels in wild brown trout (*Salmo trutta*) populations in Austrian rivers. BMC Vet Res 16:40.

56. Persson DB, Aspán A, Hysing P, Blomkvist E, Jansson E, Orsén L, Hällbom H, Axén C. 2022. Assessing the presence and spread of *Renibacterium salmoninarum* between farmed and wild fish in Sweden. J Fish Dis 45:613–621.

57. Johansen L-H, Jensen I, Mikkelsen H, Bjørn P-A, Jansen PA, Bergh Ø. 2011. Disease interaction and pathogens exchange between wild and farmed fish populations with special reference to Norway. Aquaculture 315:167–186.

58. Johansen R. 2013. Fish Health Report 2012. Norwegian Veterinary Institute Report series. Norwegian Veterinary Institute, Oslo.

59. Murray AG, Peeler EJ. 2005. A framework for understanding the potential for emerging diseases in aquaculture. Prev Vet Med 67:223–235.

60. Banner CR, Long JJ, Fryer JL, Rohovec JS. 1986. Occurrence of salmonid fish infected with *Renibacterium salmoninarum* in the Pacific Ocean. J Fish Dis 9:273–275.

61. Gutenberger S, Duimstra J, Rohovec J, Fryer J. 1997. Intracellular survival of *Renibacterium salmoninarum* in trout mononuclear phagocytes. Dis Aquat Organ 28:93–106.

62. Suzuki K, Misaka N, Mizuno S, Sasaki Y. 2017. Subclinical Infection of *Renibacterium salmoninarum* in Fry and Juveniles Chum Salmon *Oncorhynchus keta* in Hokkaido, Japan. 魚病研究 52:89–95.

63. Austin B, Rayment JN. 1985. Epizootiology of *Renibacterium salmoninarum*, the causal agent of bacterial kidney disease in salmonid fish. J Fish Dis 8:505–509.

64. Gómez FA, Tobar JA, Henríquez V, Sola M, Altamirano C, Marshall SH. 2013. Evidence of the presence of a functional Dot/Icm type IV-B secretion system in the fish bacterial pathogen *Piscirickettsia salmonis*. PloS One 8:e54934.

65. Levipan HA, Irgang R, Yáñez A, Avendaño-Herrera R. 2020. Improved understanding of biofilm development by *Piscirickettsia salmonis* reveals potential risks for the persistence and dissemination of piscirickettsiosis. Sci Rep 10:12224.

66. Zúñiga A, Aravena P, Pulgar R, Travisany D, Ortiz-Severín J, Chávez FP, Maass A, González M, Cambiazo V. 2020. Transcriptomic changes of *Piscirickettsia salmonis* during intracellular growth in a salmon macrophage-like cell line. Front Cell Infect Microbiol 9:426.

67. Gutenberger SK, Giovannoni SJ, Field KG, Fryer JL, Rohovec JS. 1991. A phylogenetic comparison of the 16S rRNA sequence of the fish pathogen, *Renibacterium salmoninarum*, to Gram-positive bacteria. FEMS Microbiol Lett 77:151–156.

68. Flores-Martin SN, Isla A, Aguilar M, Barrientos CA, Blanco JA, Rauch MC, Enríquez R, Arcos C, Yañez AJ. 2024. Optimizing culture medium to facilitate rapid *Renibacterium salmoninarum* growth. Aquaculture 587:740798.

69. Lyngstad TM, Høgåsen HR, Jansen MD, Nilsen A. 2015. Risk of disease transfer with wellboats in Norway – Technical report. Rapport 15.

70. Slette HT, Salomonsen C, Størkersen K, Tveit GM, Misund A, Lona E. 2025. Biosafety in Norwegian aquaculture—Risks and measures in RAS facilities and well-boats. Rev Aquac 17:e12979.

71. Pedersen K, Skall HF, Lassen-Nielsen AM, Nielsen TF, Henriksen NH, Olesen NJ. 2008. Surveillance of health status on eight marine rainbow trout, *Oncorhynchus mykiss* (Walbaum), farms in Denmark in 2006. J Fish Dis 31:659–667.

72. Bregnballe J. 2022. A guide to recirculation aquaculture. Food and Agriculture Organization of the United Nations, Rome, Italy. http://www.fao.org/documents/card/en/c/cc2390en. Retrieved 2 December 2024.

73. Drønen K, Roalkvam I, Nilsen H, Olsen AB, Dahle H, Wergeland H. 2022. Presence and habitats of bacterial fish pathogen relatives in a marine salmon post-smolt RAS. Aquac Rep 26:101312.

74. Noble AC, Summerfelt ST. 1996. Diseases encountered in rainbow trout cultured in recirculating systems. Annu Rev Fish Dis 6:65–92.

